# Foxh1 is a locus-specific PRC2 recruiter governing germ layer silencing

**DOI:** 10.1101/2025.09.21.677640

**Authors:** Jin Cho, Clark L. Hendrickson, Nathan Mar, Ira L. Blitz, Margaret Fish, Wenqi Wang, Ken W.Y. Cho

## Abstract

Polycomb Repressive Complex 2 (PRC2) establishes H3K27me3 marks to shape spatiotemporal gene expression during embryogenesis. While its dysregulation is linked to developmental disorders, cancer, and aging, the mechanisms guiding PRC2 to specific genomic loci remain a subject of ongoing debate. A prevailing model proposes that PRC2 recruitment occurs via its intrinsic affinity for chromatin rather than through sequence-specific transcription factors. Here, we provide evidence that the maternally deposited pioneer transcription factor Foxh1 plays a critical role in directing PRC2 to specific genomic loci during zygotic genome activation in *Xenopus*. Foxh1 is a critical transcription factor mediating Nodal signaling, but it also plays an earlier role by pre-binding enhancers prior to signaling activation. This pre-binding is essential for forming enhanceosome complexes that trigger mesendodermal gene expression and drive gastrulation, in cooperation with other maternal transcription factors. Using maternal Foxh1-null embryos, we demonstrate that Foxh1 directly recruits Ezh2, the catalytic subunit of PRC2, to Foxh1-bound loci. Loss of Foxh1 impairs Ezh2 recruitment, leading to a global reduction in H3K27me3. These findings support a dual-function model in which Foxh1 not only activates endodermal gene expression in endoderm, but also recruits PRC2 to silence the same genes in ectoderm. This dual activity of Foxh1 allows the spatially coordinated epigenetic states of the endodermal gene regulatory program during early embryogenesis.

**Highlights:**

1. Foxh1 recruits Ezh2 to deposit H3K27me3 at specific loci in early embryos
2. Maternal Foxh1 loss reduces H3K27me3 and disrupts lineage segregation
3. Foxh1 both activates and represses endoderm genes in a context-specific way

## INTRODUCTION

A central question in developmental biology is to understand how the genome is read and interpreted to orchestrate gene expression during progenitor cell differentiation. Animal cloning experiments in *Xenopus* showed that the fertilized egg can reprogram differentiated somatic nuclei, indicating a resetting of epigenetic information (Gurdon et al., 1958). During normal development, the genome of fertilized zygotes undergoes multiple rounds of cell division, while gradually reprogramming germ cell-derived chromatin into an early embryonic epigenetic landscape. The first critical transition in gene function occurs during zygotic genome activation (ZGA), when maternally deposited transcription factors (TFs) and chromatin modifiers initiate embryonic gene expression and begin to shape the embryonic epigenome (Biltz and Cho, 2021; Kojima et al., 2025). Understanding how active and repressive chromatin domains are established and organized during the extensive epigenetic remodeling of the genome remains a major question in biology.

In *Xenopus*, the embryonic genome is relatively absent of histone modifications prior to and during early ZGA, reflecting an uncommitted transcriptional state (Akkers et al., 2009; Gupta et al., 2014; Hontelez et al., 2015). As development proceeds, active marks like H3K27ac and H3K4me1 accumulate in a cell type–specific manner, often at sites bound by maternal TFs (Gupta et al., 2014; van Heeringen et al., 2014; Jansen et al., 2022: Paraiso et al., 2019). Maternally provided TFs guide histone modifiers to specific genomic loci and help establish deposition of active histone marks (Hontelez et al., 2015; Paraiso et al., 2025). In addition to depositing active marks on the genome, the deposition of repressive marks such as H3K27me3 is also critical for silencing inappropriate gene expression and preventing transcriptional misregulation. H3K27me3 deposition is controlled by Polycomb Repressive Complex 2 (PRC2), which is composed of a core tetrameric complex, Ezh2/Ezh1, Eed, Suz12, and Rbbp4/7, where Ezh2 is the key methyltransferase essential for H3K27me3 deposition (Cao and Zhang, 2004; Pasini et al, 2004). PRC2 is divided into two subtypes, PRC2.1 containing a PCL homolog, or PRC2.2 containing JARID2 and AEBP2 (Blackledge et al, 2015; Laugesen et al., 2019). PRC2.1 and PRC2.2 coordinate establishment and stabilization of H3K27me3 marks across the genome.

In *Drosophila*, Polycomb group (PcG) proteins are recruited to chromatin through specific cis-regulatory DNA sequences known as Polycomb Response Elements (PREs) (Lyko and Paro, 1999). The first PRE-binding protein to be identified was Pleiohomeotic (Pho), a sequence-specific DNA binding factor that directly mediates PcG recruitment (Klymenko et al., 2006). In contrast, although PcG proteins are also known to bind chromatin in vertebrates, no PRE-like elements have yet been identified. TFs, such as E2f, Mga, and Max, have been implicated in interactions with PRC1, but not with PRC2 (Gao et al., 2012). Mtf2 directly interacts with PRC2, but it is not a sequence-specific DNA-binding TF (Perino et al., 2018). Runt, Rest, and Snail TFs interact with PRC1 and/or PRC2, but these interactions appear to be context-specific rather than broadly required (Herranz et al., 2008 Dietrich et al., 2012; Tien et al., 2015). To date, a sequence-specific TF that directs PRC2 recruitment in early vertebrate embryos has yet to be identified.

Foxh1, a forkhead domain TF, was originally identified as a critical mediator of Nodal signaling through its interaction with Smad2/3 and Smad4, and plays a central role in mesendodermal cell differentiation (Chen et al., 1997). Its mRNA is maternally supplied in the *Xenopus* egg, and ubiquitously expressed. We previously found that Foxh1 binds to the embryonic genome prior to the deposition of histone epigenetic marks (Charney et al., 2017). To uncover the mechanistic link between Foxh1 binding and epigenetic modifications, mass spectroscopy was performed using Foxh1 as bait (Zhou et al., 2022; 2023). This analysis identified key components of the PRC2 complex including Ezh2, Suz12, Jarid2, and Rbbp7, suggesting a potential interaction between pioneer TF Foxh1 and PRC2.2.

We investigated the physical interaction and the genome-wide occupancy of Foxh1 and Ezh2 in early *Xenopus* embryos using Foxh1 mutant lines. Our results discovered that Foxh1 regulates a cohort of endodermally-expressed genes by acting as a dual-function TF. In endodermal cells, Foxh1 promotes endodermal gene activation, whereas in ectodermal cells, it facilitates their repression complex via Ezh2, leading to H3K27me3 deposition and silencing of endodermal gene expression. This dual activity of Foxh1 allows the spatially coordinated epigenetic states of the endodermal gene regulatory program during early embryogenesis.

## RESULTS

### Polycomb repressive complex 2 (PRC2) associated protein Ezh2 specifically interacts with Foxh1

The PRC2 complex is responsible for the deposition of the repressive histone mark H3K27me3 to silence target genes. While some TFs have been implicated in regulating PRC2-mediated H3K27me3 deposition, direct evidence for their locus-specific targeting in vertebrates is limited or inconclusive (Blackledge et al., 2015). Interestingly, we found that the maternally expressed pioneer TF Foxh1 physically interacts with core PRC2 components such as Suz12, Eed, RBBP4, and Ezh2 (Zhou et al., 2022). This finding raises the intriguing possibility that Foxh1 may play a role in locus-specific recruitment of PRC2 during early vertebrate embryogenesis. We first examined the interaction specificity of Foxh1. Immunoprecipitation followed by western blot analysis shows that Ezh2 forms a complex with Foxh1, but not with Foxi2, another maternally expressed Forkhead family member, or with Sox3, a maternally-expressed HMG box pioneer TF (Fig 1A). This suggests that the Ezh2-Foxh1 interaction is specific. To demonstrate their co-binding at cis-regulatory modules (CRMs) in developing embryos, we performed a sequential chromatin immunoprecipitation-quantitative PCR (ChIP-qPCR) analysis. ChIP was first performed using Ezh2 antibody, followed by a second round of ChIP using Foxh1 antibody (Fig 1B). Three enhancer regions belonging to *pitx2, gata2,* and *zic3*, are enriched over the control, demonstrating that Ezh2 and Foxh1 co-bind to these CRMs in early *Xenopus* embryos, prior to gastrulation.

**Figure 1.**
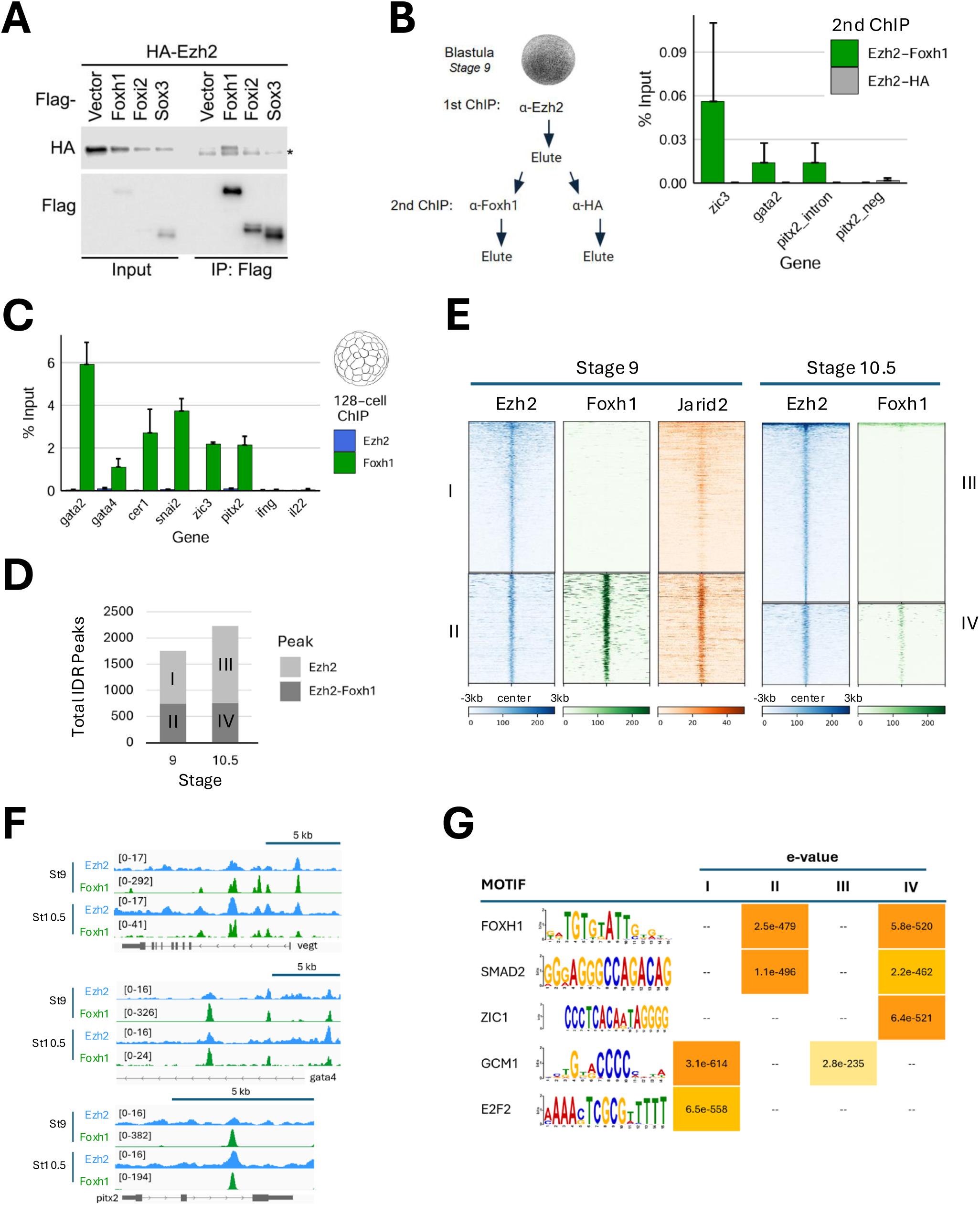
Interaction between Foxh1-Ezh2 and their occupancy at regulatory regions revealed by sequential ChIP-qPCR and ChIP-seq. A) Immunoprecipitation followed by western blotting revealed a specific interaction between Foxh1 and Ezh2. HEK293T cells were co-transfected with HA-Ezh2 and vectors expressing Foxh1, Foxi2, or Sox3 for 24 hours, and immunoprecipitation was performed using Flag-M2 beads. B) Schematic of sequential ChIP-qPCR analysis. First round ChIP for endogenously expressed Ezh2 was performed followed by elution of the chromatin, splitting of the eluate into two, which were then subjected to a second round of ChIP for endogenously expressed Foxh1, or using anti-HA as a control for background binding levels. Percent (%) Input recovery shows co-occupancy of Foxh1 and Ezh2 at enhancer regions upstream of *zic3* (5kb upstream element), *gata2* (0.7kb upstream element), and within the second *pitx2* intron. The signal from control Ezh2-HA was barely detectable. C) Bar graph showing Foxh1 and Ezh2 enrichment (% Input) at target loci at 128-cell stage. D) Number of high-confidence ChIP-seq Ezh2 peaks overlapping with Foxh1 peaks identified via Irreproducible Discovery Rate (IDR) analysis. Type I and II peaks are from stage 9 (blastula), type III and IV peaks are from stage 10.5 early (early gastrula). E) Heatmap showing Foxh1 ChIP-seq and stage 9 Jarid 2 ChIP-seq signal centered on Ezh2 IDR peak summits. Class I and III are Ezh2-only peaks, class II and IV peaks overlap with Foxh1 peaks. F) Genome browser tracks displaying co-enrichment of Ezh2 and Foxh1 at *vegt*, *gata4* and *pitx2* loci. G) Motif analysis under clustered Ezh2 peaks reveals enriched transcription factor binding motifs in each class.

### Maternal Foxh1 occupancy precedes and marks future Ezh2-bound sites

Ezh2 protein and transcripts are maternally supplied. A related gene, *ezh1*, is expressed at a very low level, and the protein is not detectable in the egg (Owens et al., 2016; Peshkin et al., 2019, Figure S1A). We determined when Ezh2 is recruited to the mesendodermal enhancers. Previously, Foxh1 was shown to bind to CRMs of mesendodermally-expressed genes by 32-cell stage and persists until the early gastrula stage (Charney et al., 2017). However, Ezh2 recruitment to the CRMs was not observed even at 128-cell stage (Fig 1C). Ezh2 ChIP-seq analysis revealed 1756 peaks at stage 9 (late blastula) and 2,231 peaks at stage 10.5 (early gastrula) (Fig 1D, Table S1). To examine the relationship between Ezh2 and Foxh1 binding, peaks were categorized based on their overlap with Foxh1. At stage 9, 740 Ezh2 peaks overlapped with Foxh1 (Class II), while 1016 peaks did not (Class I) (Fig 1D). Similarly, at stage 10.5, 757 of 2,231 Ezh2 peaks overlapped with Foxh1 (Class IV), and 1474 showed no overlap (Class III) (Fig 1D, Table S1). Genome browser views of mesendodermally-expressed genes such as *vegt, gata4,* and *pitx2* reveal multiple enhancers co-localized by Ezh2 and Foxh1. These data highlight the likelihood of both Foxh1-associated and Foxh1-independent recruitment modes of PRC2. We also aligned stage 9 ChIP-seq data for Jarid2 (van Heeringen et al., 2014), which revealed strong overlap with Class II Ezh2 peaks, suggesting that the Ezh2-Foxh1 co-bound regions are part of a functional PRC2.2 complex (Fig 1E).

To identify TFs that may mediate Ezh2 binding during early germ layer specification, a MEME-ChIP motif enrichment analysis was performed using a 500bp window surrounding the summits of Ezh2 peaks present in classes I-IV (Fig 1G). Motif analysis of class II and IV peaks revealed significant enrichment of Foxh1 and Smad2 at stages 9 and 10.5. These results suggest that a subset of Ezh2 target sites are associated with Foxh1/Smad2-regulated mesendodermally-expressed genes. Gene Ontology (GO) analysis on the nearest genes to these peaks further support this interpretation, showing stronger enrichment for terms related to endoderm formation (Fig S1B). Ezh2-bound peaks lacking Foxh1 (classes I and IV) show enrichment of GCM1 and E2F2 motifs. A temporal expression time course in *Xenopus tropicalis* shows that neither *gcm1* nor its closely related *gcm2* is expressed during early embryogenesis (Fig S1C). However, the binding motif of the zinc finger TF Zbtb10 (GNNCCCC) closely resembles the GCM1 motif. The *zbtb10* gene is maternally expressed and has been shown to function as a repressor of Sp1 TF (Mertens-Talcott et al., 2013; Fig S1D). As for *e2f2,* this gene is absent in *Xenopus*, but six other E2f family members (*e2f1, e2f3, e2f4, e2f5, e2f6,* and *e2f8*) are maternally expressed (Fig S1E). These TFs may guide PRC2 recruitment. GO analysis of genes near Class I and III peaks revealed strong enrichment for processes including cell migration involved in gastrulation, cell fate specification, and brain development (Fig S1F), suggesting Foxh1-independent modes for PRC2 in regulating cell migration and neural development.

### Generation and characterization of *foxh1 CRISPR* mutants

In order to determine the role of Foxh1 in Ezh2 recruitment during embryonic development, we generated biallelic *foxh1* mutant *Xenopus tropicalis* lines using a CRISPR/Cas9 genome editing approach (Blitz et al., 2013; Nakayama et al., 2013; Guo et al., 2014). sgRNA targeting the N-terminal region of Foxh1 was co-injected with Cas9 mRNA into 1-cell stage embryos (Fig 2A). We achieved high fidelity (>90%) in generating biallelic mutations, determined by PCR amplification of F0 embryonic genomes, followed by Sanger sequencing amplicons and deconvolution analysis (Brinkman et al., 2014; Blitz and Nakayama, 2022). Injected F0 embryos were raised to adulthood and females were further characterized for frequency of germline transmission of *foxh1* mutant alleles. We sought females showing 100% transmission of indels leading to frameshifts to create maternal *foxh1* knockout embryos by fertilizing eggs from F0 CRISPR females with wild-type (WT) sperm (Fig 2A). All eggs from an F0 female carry out-of-frame indels in *foxh1* (Fig S2A, B), and genotyping of F1 embryos confirms they are genetically heterozygous, inheriting mutant maternal alleles and wild-type paternal alleles, with no paternal transcripts present in the maternal RNA pool. F1 embryos resulting from two F0 female lines were 100% embryonic lethal, displaying severe axial defects and cyclopic phenotypes (Fig 2B). Western blot analyses of F1 embryos at pre-ZGA (stage 6) and blastula (stage 9) confirmed that Foxh1 protein is absent (Fig 2C), demonstrating that frameshift indels lead to loss of expression of maternal Foxh1 protein. To confirm that Foxh1 loss caused the phenotypes, we rescued function by injecting wild type *foxh1* mRNA into embryos from an F0 mutant female fertilized with wild type sperm (hereafter, referred to as maternal mutant *foxh1* (M*foxh1*) embryos) (Fig 2B). Transient expression from injected *foxh1* mRNA (25 pg) fully rescues normal development and these maternal loss-of-function *foxh1*^+/-^ animals are viable to adulthood (while all siblings not fully restored to wild type *foxh1* died during early embryogenesis due to defects as shown in Fig 2B). We conclude that the observed M*foxh1* embryonic phenotypes are due to the absence of maternal Foxh1 protein.

**Figure 2.**
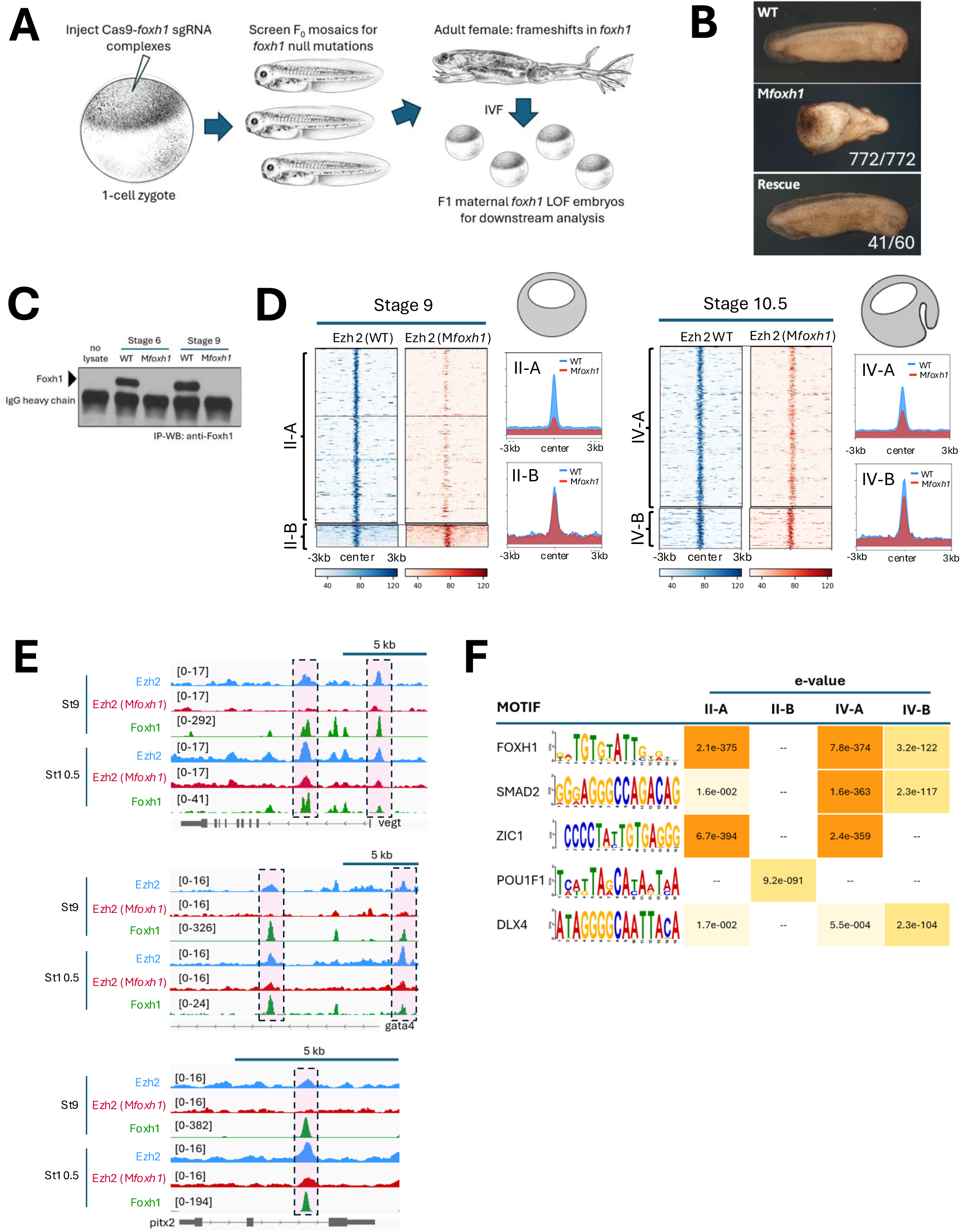
Foxh1 mutant embryos reveal locus-specific recruitment of Ezh2 to the genome. A) Schematic illustrating the generation of maternal *foxh1* deficient (Mf*oxh1*) embryos using CRISPR/Cas9-mediated mutagenesis. *foxh1* mutations were generated at high efficiency and confirmed in adult F0 females. B) Phenotypic comparison of wild type (top), M*foxh1* mutant (middle), and rescued embryos (bottom) after injection of 25 pg wild type *foxh1* mRNA. C) Western blot using anti-Foxh1 antibody confirms the absence of Foxh1 protein in M*foxh1* embryos at stages 6 and 9. D) Ezh2 ChIP-seq analysis at class II and IV Foxh1-Ezh2 co-bound regions in stage 9 and 10.5 embryos. Peaks in subsets II-A and IV-A show reduced Ezh2 binding in M*foxh1* embryos, whereas II-B and IV-B peaks remain largely unaffected. E) Genome browser view of Ezh2 peaks at *vegt*, *gata4,* and *pitx2* loci in wild-type and M*foxh1* embryos. Boxed regions highlight Ezh2 peaks that are specifically diminished in M*foxh1* embryos. F) Motif enrichment analysis of class II-A, II-B, IV-A and IV-B peaks identified transcription factor binding motifs with Foxh1-dependent or -independent Ezh2 peaks. *Xenopus* illustrations © Natalya Zahn (2022).

### Foxh1 is required for Ezh2 recruitment to specific loci

To determine whether Foxh1 is required for Ezh2 recruitment to the embryonic genome, Ezh2 ChIP-seq was performed in wild type and M*foxh1* mutant embryos. ChIP-seq data of stage 9 and 10.5 M*foxh1* embryos revealed that 91% (673 out of 740 peaks, class II-A) of stage 9, and 82% (623 peaks out of 757, class IV-A) of stage 10.5 Ezh2-bound peaks are Foxh1 dependent (Fig 2D, Table S1). Genome browser view shows that Ezh2 peaks on enhancers of endodermally-expressed genes *vegt*, *gata4,* and *pitx2* are significantly reduced or completely absent in M*foxh1* embryos (Fig 2E). The results show that Foxh1 mediates a significant portion of Ezh2 binding, with the rest likely guided by other maternal TFs. Binding site enrichment analysis on the regions contained within these Ezh2 peaks revealed Foxh1 motifs along with Smad2 motifs being highly enriched in Foxh1-dependent Ezh2 peaks (Fig 2F, class II-A and IV-A). This suggests that these target genes are regulated by Nodal signaling, and represent endodermally-expressed genes. Other TF motifs were also discovered in this enrichment analysis, including those for Zic TFs. While Zic 1 is not expressed maternally, Zic2 is expressed maternally in *Xenopus* (Fig S2C), and its loss of function results in gastrulation defects and abnormalities in anteroposterior axis formation (Houston & Wylie, 2005). In contrast, Foxh1-independent Ezh2 peaks were enriched for Pou motifs at stage 9 and Dlx motifs at stage 10.5, suggesting alternative recruiting mechanisms. Together, these findings support a model in which Ezh2 is recruited to the genome by multiple maternal TFs.

### Foxh1 controls spatial H3K27me3 deposition to restrict endodermal gene expression to the appropriate germ layer

H3K27me3 deposition by PRC2 appears during the blastula stage and increases by early gastrulation (Akkers et al., 2009; Gupta et al, 2014; van Heeringen et al., 2014), coinciding with onset of the germ layer specification and gastrulation. In order to determine whether the H3K27me3 landscape is regionally patterned to reflect germ layer specific gene expression, we first identified genes with transcripts enriched in ectoderm and endoderm of early gastrula (st 10.5) embryos using RNA-seq of dissected tissue fragments (Blitz et al, 2017) (Fig 3A, Table S2). We then mapped Ezh2 peaks to the nearest genes within 20kb and compared these genes to the RNA-transcripts from dissected early gastrula tissue fragments to determine whether they were ectodermally or endodermally localized (Fig 3B). This analysis revealed 86 ectodermally expressed genes are co-localized with Ezh2 and Foxh1 peaks at the regulatory regions, as well as 219 endodermally expressed genes, suggesting Ezh2 and Foxh1 preferentially regulate endodermally expressed genes (Fig 3B, Table S2). To assess germ layer specific H3K27me3 patterns, we performed ChIP-seq analysis of ectoderm (animal) and endoderm (vegetal) explants from wild-type and M*foxh1* mutant embryos. Mesoderm was excluded from analysis due to the risk of contamination from neighboring germ layers during manual dissection. We compared H3K27me3 levels in both ectoderm and endoderm explants dissected from wild type and M*foxh1* embryos (Fig 3C, D, Table S3). In both ectoderm and endoderm explants, H3K27me3 levels on ectodermally-expressed genes were low in wild type and M*foxh1* embryos (Fig 3C, left panels, Fig S3). The average read density (PRKM) across the 20kb upstream, the entire gene body, and the 20kb downstream region of the genes remains consistently below 10. These results indicate that H3K27me3 is not significantly deposited at ectodermally-expressed gene loci during early gastrulation.

**Figure 3.**
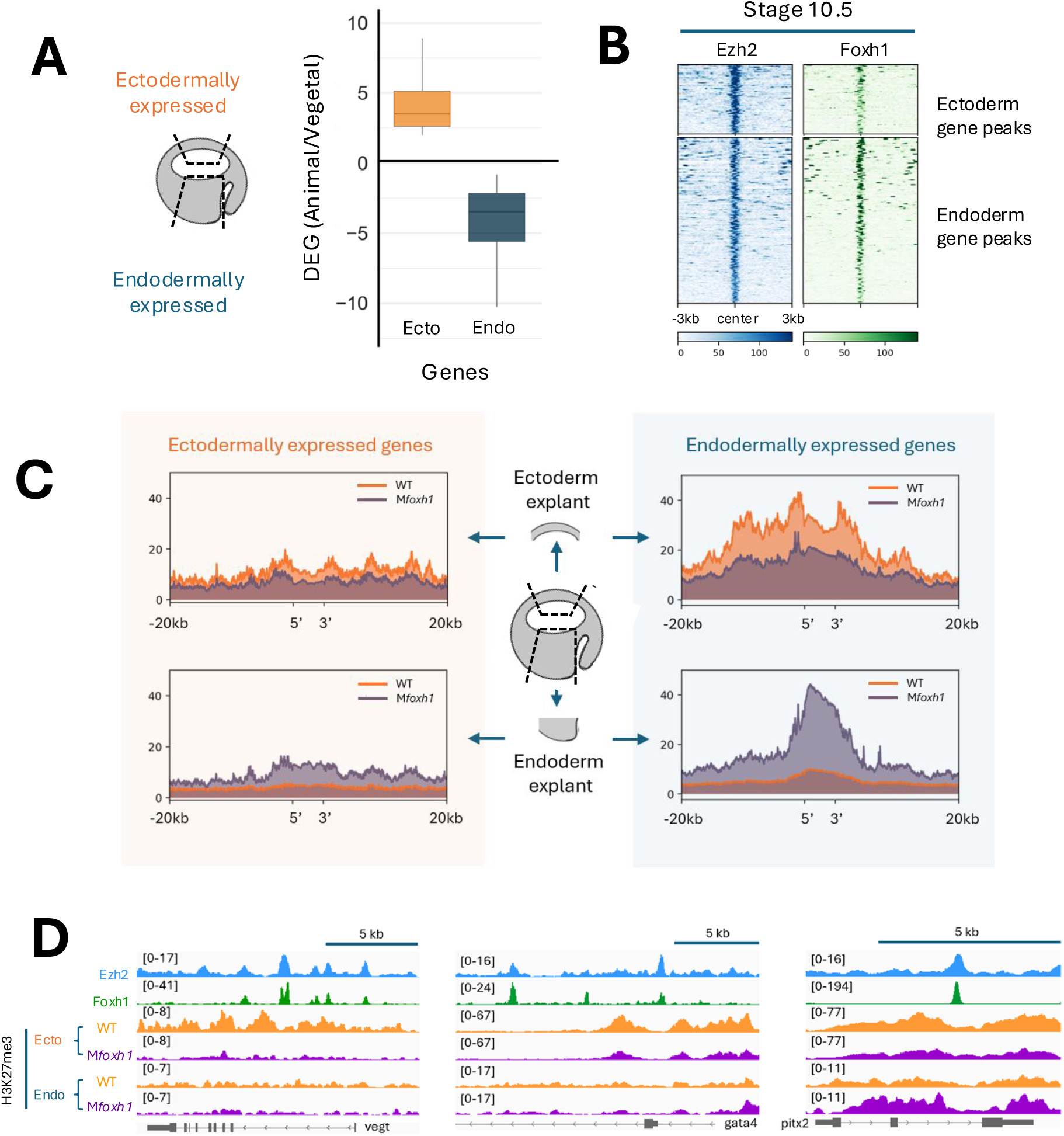
Loss of Foxh1 alters H3K27me3 deposition at endodermal gene loci. A) Identification of genes enriched in the ectodermal and endoderm germ layers of early gastrula embryos. B) Ezh2 and Foxh1 peaks were mapped to germ layer specific genes, with Ezh2 peak summits centered to assess their binding patterns at ectoderm- and endoderm-expressed genes. C) H3K27me3 deposition profiles for ectodermally and endodermally expressed genes, measured in isolated ectoderm and endoderm explants from wild-type and M*foxh1* embryos. D) Genome browser view showing Ezh2, Foxh1, and H3K27me3 profiles across different germ layers in wild-type and M*foxh1* embryos at *vegt*, *gata4,* and *pitx2 loci*.

We also examined the H3K27me3 profiles of endodermally-expressed genes such as *vegt, gata4,* and *pitx2* in both ectoderm and endoderm explants (Fig 3D, Fig S3, Table S3). These genes largely lack the H3K27me3 repressive mark in the endodermal explants (Fig 3D), consistent with their absence of transcripts in the region. In contrast, wild type ectodermal explants show high levels of H3K27me3 at these loci, indicating strong repression of endodermally-expressed genes in the ectoderm (Fig 3C, D), similar to previous findings (Akker et al., 2009). However, in M*foxh1* mutant embryos, we observed a significant reduction in H3K27me3 deposition at endodermally-expressed gene loci in ectoderm explants, indicating that Foxh1 is required for proper PRC2 recruitment and H3K27me3 marking of these regions (Fig 3C right top, Fig 3D, Fig S3). This is consistent with the model that Foxh1 facilitates locus-specific recruitment of PRC2 via Ezh2. Interestingly, in the endoderm explants of M*foxh1* embryos, overall H3K27me3 levels were significantly elevated. This unexpected result suggests that, in the absence of Foxh1, enhancer regions of some endodermally-expressed genes become vulnerable to constitutive PRC2 recruitment, leading to H3K27me3 deposition at regulatory elements that are typically active.

## DISCUSSION

Here we propose a model for the regulation of H3K27me3 deposition on embryonic chromatin (Fig 4). Maternal Foxh1 is ubiquitously expressed in early embryos (Chiu et al., 2014; Blitz et al., 2017; Paraiso et al., 2019), and binds to endodermally-expressed gene enhancers during the early cleavage stages, regardless of whether cells are located animally or vegetally (left). We previously showed that localized endodermal TFs such as VegT and Otx1 form enhanceosome complexes with Foxh1 at endodermally-expressed gene loci in the endoderm (Paraiso et al., 2019). After onset of ZGA (middle), Nodal signaling becomes active in the endoderm, leading to the recruitment of Smad2/3 and Smad4 at the Foxh1-bound endoderm enhancers. In contrast, in ectodermal cells (left), Foxh1 remains bound to endodermally-expressed gene enhancers, but fails to form enhanceosome complexes due to the absence of localized maternal endodermal TFs. During ZGA, which coincides with the mid-blastula stage, in the ectodermal region, Ezh2 is recruited to Foxh1 bound sites to promote loci-specific assembly of the PRC2 complex (middle). This results in the deposition of H3K27me3 and the establishment of an active repressive state of the ectodermal genes. It remains unknown why Ezh2 recruitment does not occur during the cleavage stages and is instead delayed until the blastula stage. Several possibilities may explain this delay, including the gradual accumulation of Ezh2 to a threshold level required for the PRC2 complex formation, the absence or limited stoichiometry of essential PRC2 components, or delayed nuclear localization of Ezh2 due to cytoplasmic retention mechanisms.

**Figure 4:**
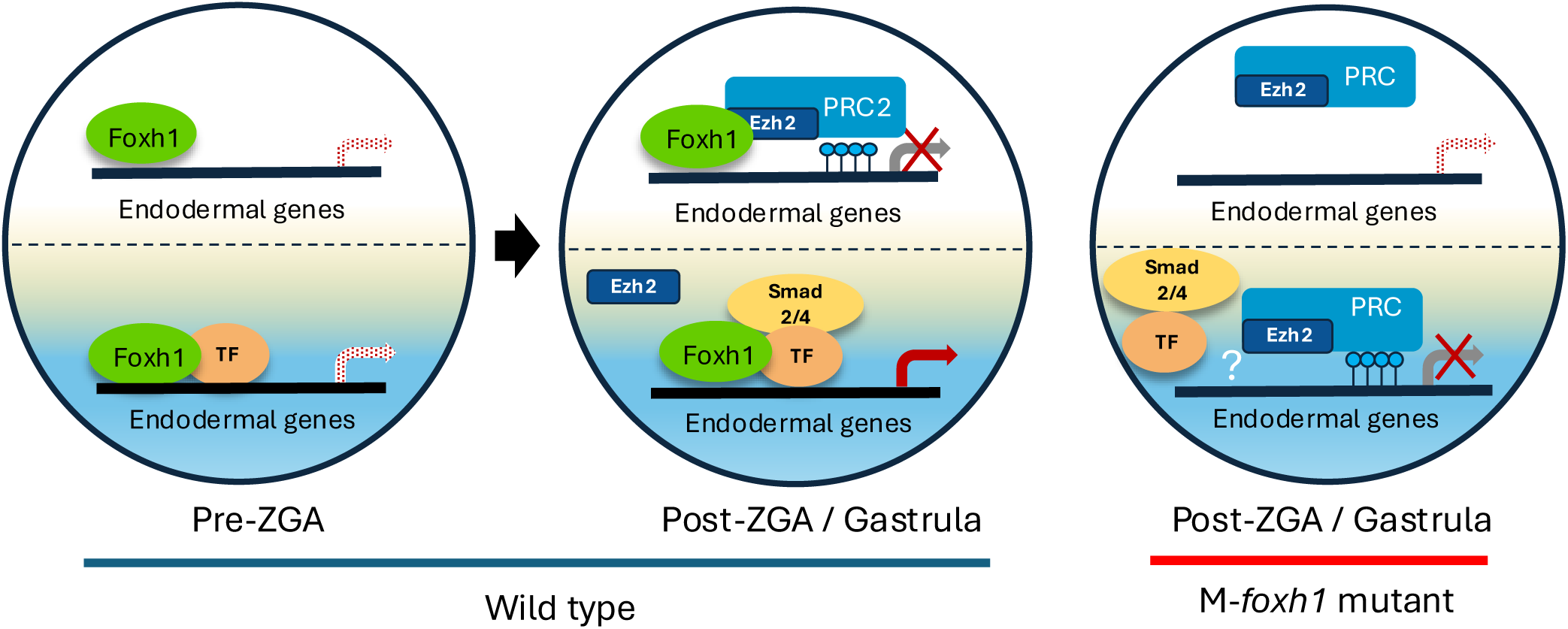
Model for Foxh1-mediated regulation of H3K27me3 deposition during early development. The left embryo represents pre-ZGA and post-ZGA/early gastrula stages in wild-type embryos. The right embryo represents an M*foxh1* embryo. In wild-type embryos at pre-ZGA stages, Foxh1 is ubiquitously expressed and binds to endodermal gene enhancers in both animal (prospective ectoderm) and vegetal (endodermal) cells. These enhancers are also occupied by other endoderm-localized maternal TF(s) such as Vegt, which prevent Ezh2/PRC2 recruitment to endodermally-active loci in the vegetal hemisphere. During post-ZGA/gastrula stages, Nodal signaling in the endoderm recruits Smad2/4 to the enhancers, activating endodermally-expressed genes. In contrast, within the ectoderm, the Ezh2-PRC2 complex is recruited to endodermal enhancers, where it deposits H3K27me3 marks to repress their expression. In M*foxh1* embryos, loss of Foxh1 impairs Ezh2-PRC2 recruitment to enhancers of endodermally-expressed genes in the ectoderm, preventing their repression. In contrast, in the endoderm, loss of Foxh1-mediated enhanceosome assembly renders these enhancers susceptible to H3K27me3 deposition, possibly through a default silencing mechanism.

The model in Figure 4 is attractive because it explains how endodermal gene expression is restricted to the appropriate cell lineage and highlights the dual, opposing roles of Foxh1 across the primary germ layers. We propose that during normal development, Foxh1-dependent enhanceosomes assembled prior to ZGA play a critical role in protecting endodermal enhancers from PRC2-mediated repression. Conversely, Foxh1-dependent recruitment of PRC2 in the ectoderm ensures active repression of these genes, potentially restricting the endodermal expression program to the endoderm. Therefore, PRC2 can be locally and selectively recruited to specific genomic loci through Foxh1 pioneer transcription factor, while PRC2 also constitutively surveys the genome to maintain repressive states. Since some Ezh2-bound regions are not co-occupied by Foxh1 but contain motifs for other transcription factors (Fig 1E, G), additional maternally expressed TFs may similarly recruit PRC2 in a non-overlapping, locus-specific manner.

One surprising observation of *foxh1* mutant embryos is that some endodermally-expressed genes become marked by H3K27me3 in endoderm, whereas in the wild type embryo this is not observed (Fig 3C, right bottom). While the underlying mechanism of this ectopic H3K27me3 is unclear, a possible explanation may be that in the absence of Foxh1, enhanceosome complexes fail to assemble at endodermal enhancers. This disruption likely results in exposing these enhancer regions, which are then recognized by PRC2. Perhaps, PRC2 complexes continuously scan regulatory elements throughout the chromatin, and respond to the absence of transcriptional activation by depositing H3K27me3 (Hosogane et al., 2013).

Mutations in core PRC2 components have revealed their essential roles in early embryonic development. Loss-of-function mutations in genes encoding subunits such as EED or EZH2 result in embryonic lethality in mice, with development arrest at peri-implantation or during gastrulation (Schumacher et al., 1996; O’Carroll et al., 2001). In mouse ES cells, inhibition of PRC2 activity prior to or during directed 2D endoderm differentiation results in a global reduction in H3K27me3, and an increase in cells expressing both HHEX and GSC, markers of mesendoderm identity (Hölzenspies et al., 2025). These findings support the idea that PRC2-mediated repression of endoderm-specific genes is crucial for proper gene regulation. The observation that PRC2 subunit mutations in mice disrupt early germ layer specification and gastrulation is consistent with findings in *Xenopus*, where Foxh1-dependent PRC2 recruitment during early gastrulation disrupts H3K27me3 deposition at endodermal genes. In the absence of Foxh1, endodermally expressed genes are erroneously marked with H3K27me3 within the endoderm, while failing to be repressed in non-endodermal tissues. The similarity between PRC2 loss-of-function phenotypes in mice and ES cells, and Foxh1-dependent PRC2 recruitment in *Xenopus*, raises the possibility that transcription factor-mediated recruitment of PRC2 to repress endodermally expressed genes may represent a conserved regulatory mechanism in vertebrates.

## Acknowledgements

This research was funded by NIH grants R35 GM139617 and R21 HD109696 awarded to K.W.Y.C. This work utilized resources of the UCI Genomics Research and Technology Hub (GRT Hub) parts of which are supported by NIH grants to the Comprehensive Cancer Center (P30CA-062203) and the UCI Skin Biology Resource Based Center (P30AR075047) at the University of California, Irvine, as well as to the GRT Hub for instrumentation (1S10OD010794-01and 1S10OD021718-01). We thank Xenbase for providing genomic and community resources (RRID: SCR_003280), the National Xenopus Resource (NXR) (RRID:SCR_013731) for supplying *Xenopus tropicalis*, and the University of California, Irvine High Performance Computing Cluster for computational resources and support.

## Author Contributions

JC, ILB and KWYC designed the project. JC and MF performed the experiments (with the assistance of ILB), and CLH, NB and JC performed the bioinformatic analyses. JC, CLH and KWYC drafted the manuscript, and CH, ILB, JC and KWYC performed manuscript edits.

## Declaration of interests

The authors declare no competing interests

## MATERIALS AND METHODS

### Animal husbandry and embryo manipulation

*Xenopus tropicalis* adult frogs were raised and maintained under guidance from the University of California, Irvine, Institutional Animal Care Use Committee. *X. tropicalis* females were primed with 10 units (U) of human chorionic gonadotropin (Chorulon HCG, Merck and Co.) 1-3 days before use and then were boosted with 100U of HCG on the day of egg collection. Eggs were “squeezed” into BSA (1 mg/ml in 1xMMR)-coated dishes and *in vitro* fertilized using testis macerated in 1X MMR containing BSA. Sperm suspension was added to eggs, diluted with 1/9X MMR to activate the sperm, and 10 minutes after sperm addition, the fertilized eggs were de-jellied using 3% cysteine in 1/9x MMR, pH 7.8. De-jellied embryos were washed and cultured at 25°C in 1/9X MMR until the desired stage according to Nieuwkoop-Faber developmental table (Nieuwkoop and Faber, 1994).

### Co-immunoprecipitation and Western blot assays

To test the interaction between Ezh2 and Foxh1, Foxi2 and Sox3, the coding sequence of Ezh2 was cloned into pCS2+ after reverse-transcribed cDNA to generate HA-tagged fusion proteins. Foxh1, Foxi2 and Sox3 coding regions were cloned in pCS2+ for the expression of *N*-terminal 3XFLAG-tagged fusion proteins. HEK293T cells were cultured in Dulbecco’s Modified Eagle Medium (DMEM) supplemented with 10% fetal bovine serum (FBS) (R&D, S11195) and penicillin-streptomycin. HEK293T cells were transfected with various combinations of constructs using the polyethylenimine (Polysciences) method (Longo et al., 2013). Cells were directly harvested in 1X SDS loading buffer (50mM Tris-HCl pH 6.8, 2% SDS, 10% glycerol, 1% b-mercaptoethanol, 0.02% bromophenol blue), boiled at 95°C for 10 mins, and subjected to western blotting using anti-FLAG (Millipore-Sigma, F3165) and anti-HA (Millipore-Sigma, 11583816001) antibodies.

Embryonic extracts were isolated from wild type *Xenopus tropicalis* and *foxh1* mutant embryos, and subject to SDS-PAGE gel electrophoresis, followed by western blot analysis using anti-Foxh1 antibody (Chiu et al., 2014).

### CRISPR/Cas9 mutagenesis

CRISPR-mediated mutagenesis in *Xenopus* was performed as described (Blitz et al., 2013). Foxh1 sgRNA was designed to mutagenize downstream of the translation start site of *foxh1*. The *foxh1*-targeted sequence was 5’-GGCCGCCCTTGTACCGAGAG<u>GGG</u>-3′ (underline indicates the PAM sequence). Linearized pCasX plasmid (Blitz et al., 2013) used for the transcription of Cas9 mRNA using the T7 mMessage mMachine kit (Ambion). The dose range injected was 3–4 ng Cas9 mRNA/embryo and 150-200 pg *foxh1* sgRNA/embryo. The cocktail of Cas9 mRNA and *foxh1* sgRNA in 4 nL of injection volume was microinjected into a single site in the animal pole at the 1-cell stage. During microinjection, embryos were in agarose-coated plates containing 1X MMR at room temperature and then embryos were cultured in agarose-coated plates in 1/9X MMR at 24–25°C until desired stages. We also performed a Foxh1 mutant rescue experiment by injecting a total of 25 pg of wild type *foxh1* mRNA into four sites to ensure even distribution. Embryos were then allowed to develop.

### Genomic PCR and Sequencing for F1 genotyping

Individual F1 embryos were obtained by *in vitro* fertilization of eggs from Foxh1 CRISPR F0 female or wild type (WT) female with sperm from a WT male. Individual embryos were transferred to 0.2-mL PCR tubes and homogenized in 100 μl of lysis buffer (50 mM Tris, pH 8.8, 1 mM EDTA, 0.5% Tween 20, 200 μg/ml proteinase K). Embryos are incubated at 56°C for 2 h to overnight, followed by incubation at 95°C for 10 min to inactivate proteinase K. Lysates were centrifuged for 1 min at 4°C and 1 μl aliquots were used directly in 20μL PCR reactions. To assess mutagenesis efficiency, a fragment containing the CRISPR site in *foxh1* was amplified by PCR with the following flanking PCR primers: Forward: 5’-CCACTTGCTGAAGGTTCGTT-3’ and Reverse: 5’-AATATGGTGGCTTGGCGTAG-3’. PCR product was purified and eluted in 20μL and subjected to Sanger sequencing from one end. Sequencing files were then subjected to deconvolution analysis to determine the efficiency of mutagenesis (Brinkman et al. 2014; Blitz and Nakayama, 2022).

### Chromatin immunoprecipitation (ChIP) assay

ChIP using *Xenopus tropicalis* embryos was performed as previously described (Chiu et al., 2014; Charney et al., 2017). The following antibodies were used: anti-Ezh2 (Abcam ab191250), anti-Foxh1 (Chiu et al., 2014), anti-HA (Abcam ab9110), and anti-H3K27me3 (Upstate/Millipore 07-449).

For sequential ChIP assay, the first round of ChIP was performed using Foxh1 antibody. After the immunoprecipitation with the first antibody, chromatin was eluted in 1x TE, pH 8.0, with 10mM DTT, 500 mM NaCl and 0.1% SDS at 37°C for 30 minutes. The eluate was then diluted 10-fold with RIPA buffer, split into two aliquots, and then incubated with the second antibody (anti-Ezh2 or anti-HA control) and the rest of the procedure was performed as in the first round of ChIP.

### Quantitative PCR

Quantitative ChIP (ChIP-qPCR) was performed using a Roche LightCycler 480 II and SYBR Green I master mix (Roche). The following primers were used for qPCR. pitx2 intron2:

F: 5’-ATCTGCTCCCATCTCTCCAA-3’; R: 5’-CAAACAGGGCTCATTGAGGA-3’

gata2 0.7kb upstream:

F: 5’-GTCGCTCTGCTCAGCTCTTC-3’; R: 5’-CCGTTTCACAGATGTGGACT-3’

zic3 5kb upstream:

F: 5’-ggaaatggaactggggaaag-3’; R: 5’-GGGTGATCTGAGCCAAATTC-3’

pitx2 negative 20kb downstream:

F: 5’-TGCACCTAGGTTTGGGTAGG-3’; R: 5’-GAGGGTGGAAAGGGGTTAAG-3’

### Library prep

ChIP-seq libraries were generated using Nextflex ChIP-seq kit (Bioo Scientific). The quality of the libraries was measured using an Agilent Bioanalyzer 2100 and KAPA qPCR and the libraries were sequenced using an Illumina platform at the UC Irvine Genomics and Research Technology Hub.

### Sequence Alignment and Visualization

**ChIP-seq:** ChIP-seq read files were aligned to the *Xenopus tropicalis* v10.0 genome (Xenbase, https://www.xenbase.org/xenbase/static-xenbase/ftpDatafiles.jsp) using Bowtie2 v2.5.1 (Langmead et al., 2012). File conversions between .SAM, .BAM, and .BED were performed using Samtools v1.15.1 and Bedtools2 v2.30 (Li et al., 2009). Duplicate reads were removed using a Samtools pipeline: “sort -n”, “fixmate -m”, “sort”, and “markdup -r”. Reads close to mitochondrial genes were removed. **Macs2 peak-calling and IDR:** Peaks were called against a stage-matched input (Charney et al., 2017) using Macs2 v2.2.7.1 by executing the default “callpeak” options with a p-value of 0.001 (Zhang et al., 2008). Peak consistency was evaluated using Irreproducibility Discovery Rate (IDR) scripts (Li et al., 2011) against a p-value threshold of 0.01 for true biological replicates and 0.02 for pseudo replicates as specified in (Charney et al., 2017). **Profile plot generation:** Bigwig file generation was performed using deepTools v3.5.1 “bamCoverage”. Samples were normalized by read count and RPKM. deepTools was used to perform the “computeMatrix” command with the “reference-point --referencePoint ‘center’” option for IDR peak regions files and the “scale-regions” option for gene regions files. Profile plots were generated using deepTools “plotProfile”. **Box plot generation:** Average BigWig scores were calculated using deepTools “multiBigwigSummary” after splitting gene regions into ‘Upstream’, ‘Gene Body’, and ‘Downstream’ in Jupyter Notebook. Box plots were created using Matplotlib v3.10.0. **Heatmap generation:** The deepTools (Ramírez et al., 2016) “computeMatrix” command with the “reference-point - -referencePoint ‘center’” option for IDR peak regions files and the “scale-regions” option for gene regions files was also used to prepare heatmap generation. deepTools “plotHeatmap” was used to create the heatmaps, and clustering was performed by specifying the option “--kmeans 3”. **Genome browser visualization:** HOMER (V4.11) (Heinz et al., 2010) commands “makeTagDirectory” and “makeUCSCfile” were used to generate a bedgraph file. Bedgraph files were converted to .TDF files using igvtools “toTDF” in the Integrative Genomics Viewer (IGV). Sample tracks were visualized in IGV. **Gene localization analysis:** Animal (ectoderm) and vegetal (endoderm) Locally-expressed genes were identified using EdgeR to calculate differentially expressed genes between tissue dissections (Blitz et al., 2017). Ectoderm-localized and endoderm-localized genes were selected by a Log2 Fold-Change threshold of at least 1 and a False Discovery Rate (FDR) below 0.05. Next, “bedtools closest” was used to generate a list of genes within 20kb of the sample IDR peaks. The gene lists were compared to ectoderm-localized and endoderm-localized genes to determine tissue-dissected epigenetic build-up around spatially expressed genes.

### Motif Analysis

MEME-ChIP from the MEME Suite 5.5.8 (http://meme-suite.org/tools/meme-chip) was used for motif analysis (Bailey et al., 2009) with the default setting. The 500bp centered on the summit of a peak was generated as the input FASTA sequence using Bedtools2 v2.30 (Quinlan and Hall, 2010).

### The accession of published RNA-seq and ChIP-seq datasets

Published datasets used in this research can be downloaded from NCBI GEO (https://www.ncbi.nlm.nih.gov/geo/) using the GEO accession numbers: GSE85273 for stage 8,9, and 10.5 Foxh1 ChIP-seq data (Charney et al., 2017);GSE53652 for stage 10.5 Foxh1 ChIP-seq data and input DNA (Chiu et al., 2014); GSE41161 for stage 9 Jarid2 (van Heeringen et al., 2014); GSE67974 for stage 9 and 10.5 Ep300 (Hontelez et al., 2015); GSE56000 for stage 8,9, and 10.5 H3K27ac (Gupta et al., 2014); and GSE81458 for stage 10.5 tissue dissected RNA-seq data (Blitz et al., 2017).

## Supplementary Figure Legends

**Supplementary Figure 1:**
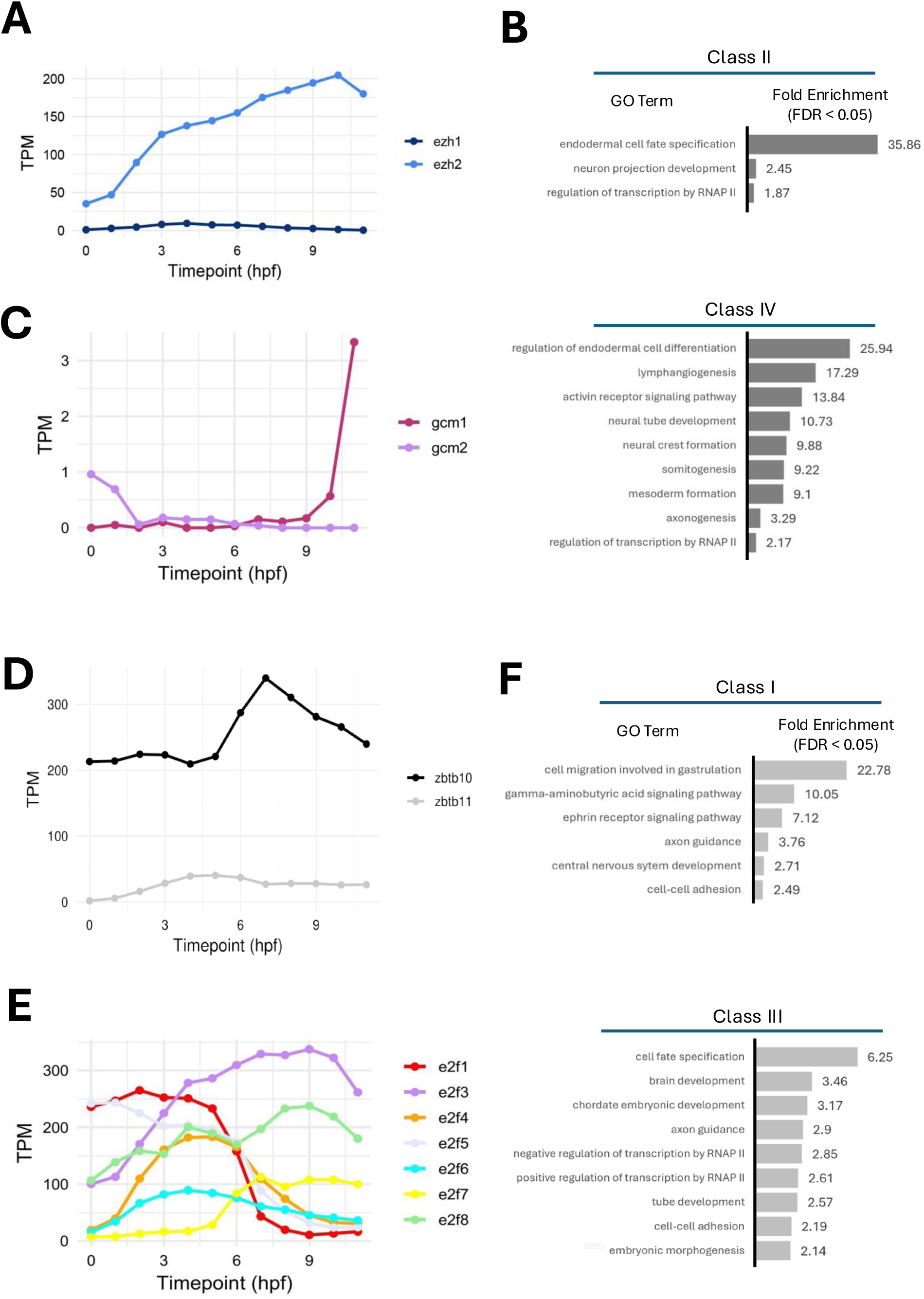
Temporal expression patterns of candidate transcription factors involved in Ezh2 recruitment to enhancers, and GG term enrichment of genes potentially regulated by Ezh2. (A). Temporal expression profiles of *ezh1* and *ezh2*. (B) GO term analysis of genes associated with Foxh1 and Ezh2 binding. (C-E) Temporal expression profiles of *gcm1/gcm2*, *zrb10/zrb11*, and the *e2f* gene family. (F) GO term analysis of genes associated with Foxh1-independent Ezh2 peaks.

**Supplementary Figure 2:**
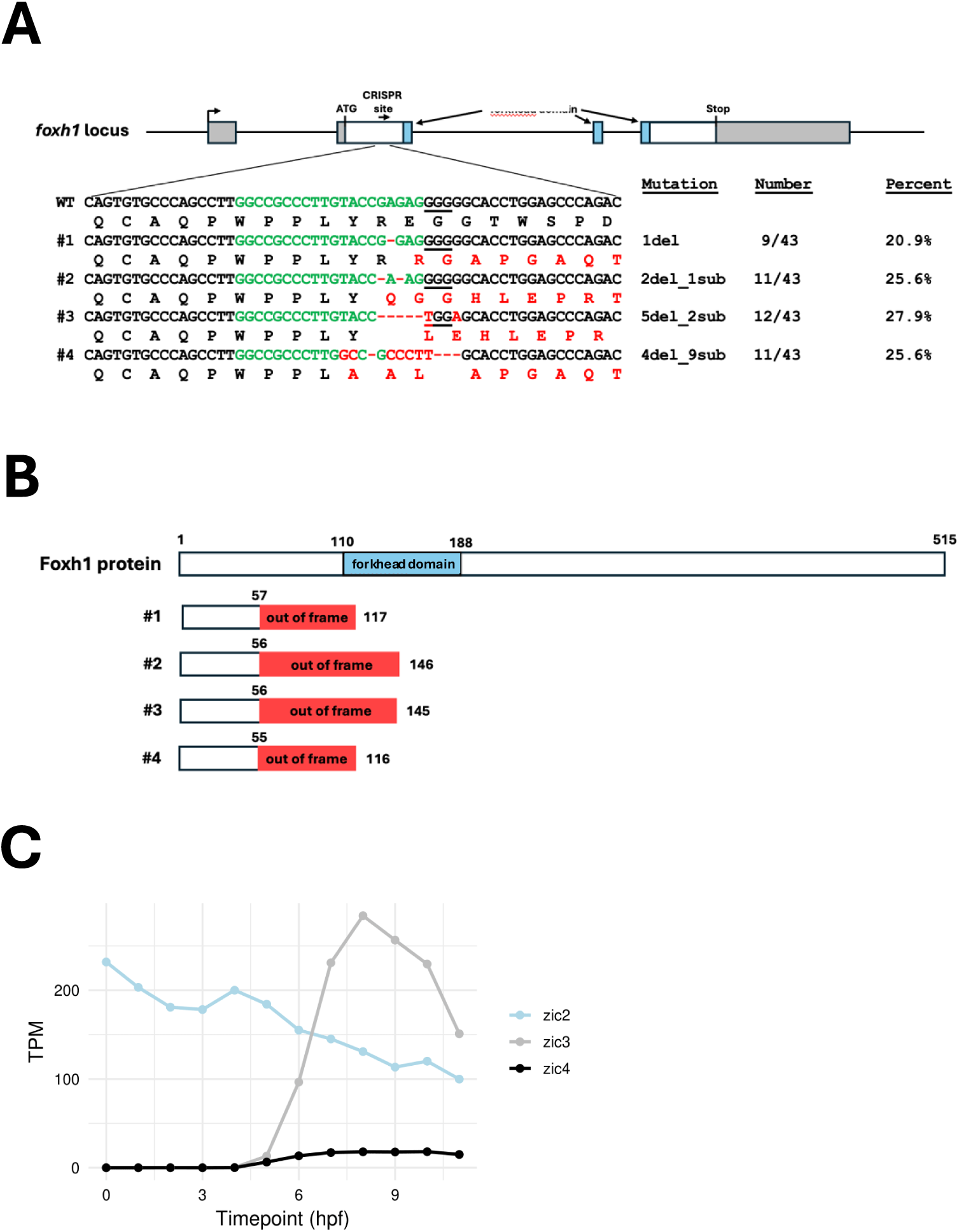
Location of indel mutations generated by CRISPR/Cas9 mutagenesis. A) Genomic PCR and Sanger sequencing used for F1 genotyping. DNA from individual F1 embryos, generated by crossing a Foxh1 CRISPR F0 female with a wild type male, was amplified by PCR and sequenced. The germline of CRISPR F0 female carried 4 types of indel mutations, with no wild type *foxh1* sequences detected, indicating that nearly all germ line cells harbored these mutations. (B) Predicted mutant Foxh1 proteins resulting from the indels identified in (A). (C) Temporal gene expression profiles of *zic* family genes.

**Supplementary Figure 3:**
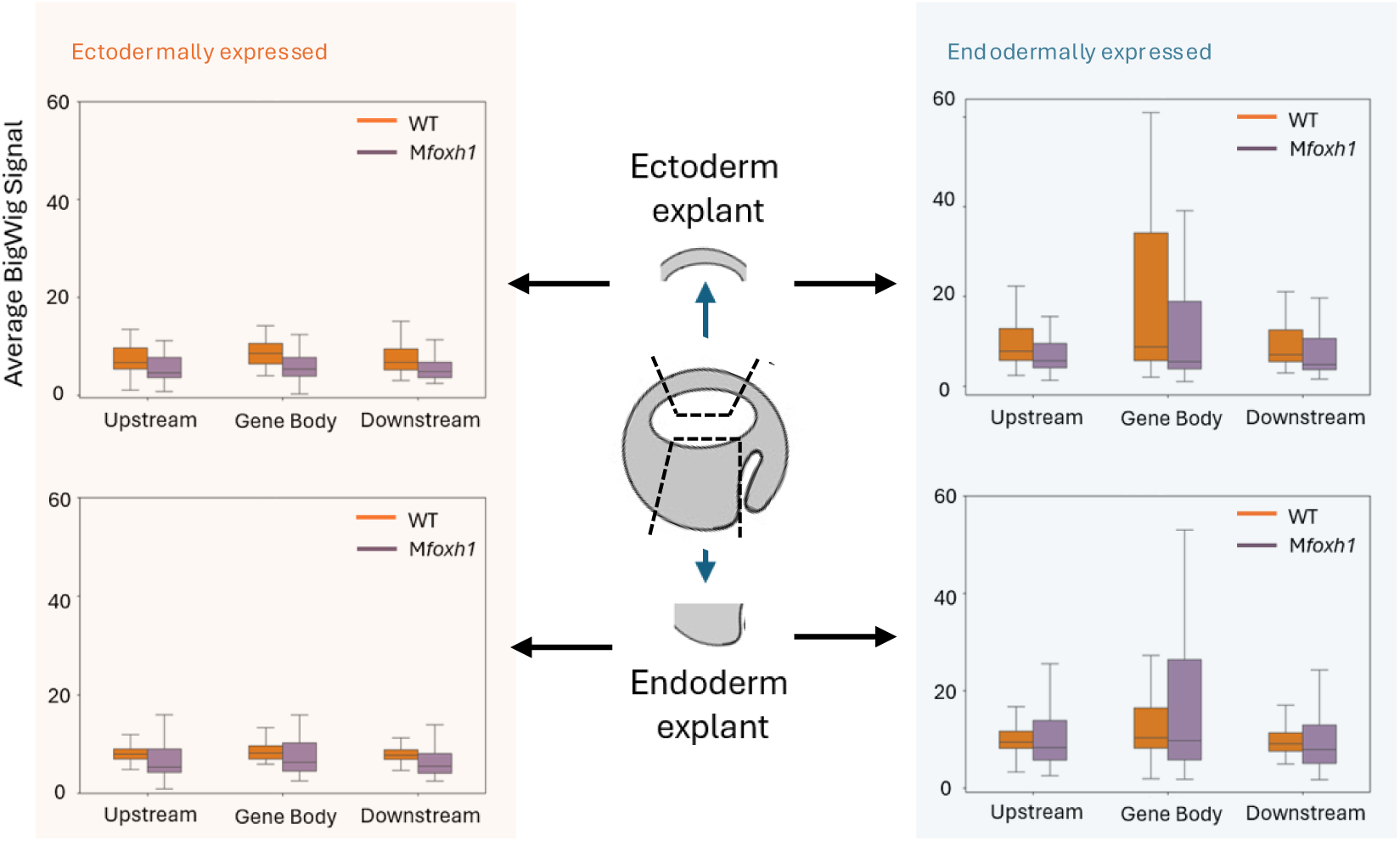
Quantification of H3K27me3 deposition profiles at ectodermally and endodermally expressed genes, measured in isolated ectoderm and endoderm explants from wild-type and M*foxh1* embryos. ‘Upstream,’ ‘gene body,’ and ‘downstream’ correspond to the 20 kb regions upstream of the TSS, across the gene body, and 20 kb downstream of the TES, respectively. Values shown represent average RPKM values over the designated region.

## Notes

### Competing Interest Statement

The authors have declared no competing interest.

